# Testing the phylogenetic gambit: how much functional diversity can we reliably conserve if we prioritize phylogenetic diversity?

**DOI:** 10.1101/243923

**Authors:** Florent Mazel, Matthew W. Pennell, Marc Cadotte, Sandra Diaz, Giulio Valentino Dalla Riva, Richard Grenyer, Fabien Leprieur, Arne O. Mooers, David Mouillot, Caroline M. Tucker, William D. Pearse

**Affiliations:** Department of Biological Sciences, Simon Fraser University, Burnaby, BC, Canada; Department of Botany, University of British Columbia, Vancouver, BC V6T 1Z4, Canada; Biodiversity Research Centre, University of British Columbia, Vancouver, BC V6T 1Z4, Canada; Department of Zoology, University of British Columbia, Vancouver, BC V6T 1Z4, Canada; Biological Sciences, University of Toronto-Scarborough, Scarborough M1C 1A4, Canada; Ecology and Evolutionary Biology, University of Toronto, Toronto, ON, Canada; Instituto Multidisciplinario de Biología Vegetal. FECFyN - Universidad Nacional de Córdoba. Casilla de Correo 495, 5000 Córdoba. Argentina; Department of Statistics, University of British Columbia, Vancouver, BC V6T 1Z4, Canada; School of Geography and the Environment, University of Oxford, Oxford OX1 3QY, U.K.; Marine Biodiversity, Exploitation, and Conservation (MARBEC), UMR 9190, Université Montpellier, Montpellier, France; Australian Research Council Centre of Excellence for Coral Reef Studies, James Cook University, Townsville, QLD, Australia; Department of Biology, University of North Carolina-Chapel Hill; Ecology Center and Department of Biology, Utah State University, Logan, Utah

**Keywords:** conservation, EDGE, evolutionary distinct, trait diversity, evolutionary diversity

## Abstract

In the face of the biodiversity crisis, it is argued that we should prioritize species in order to capture high functional diversity (FD). Because species traits often reflect shared evolutionary history, many researchers have advocated for a “phylogenetic gambit”: maximizing phylogenetic diversity (PD) should indirectly capture FD. For the first time, we empirically test this gambit using data from >15,000 vertebrate species and ecologically-relevant traits. Maximizing PD results in an average gain of 18% of FD relative to random choice. However, this average gain hides the fact that in over 1/3 of the comparisons, maximum PD sets contain less FD than randomly chosen sets of species. These results suggest that, while maximizing PD protection can help to protect FD, it represents a risky strategy.

**Statement of authorship:** FM, MP, MC, SD, GVDR, RG, AOM, CT and WP conceived the design of the study. FM and GVDR conducted the analysis. FM, RG, MP and WP interpreted the results and wrote the first draft of the manuscript. All authors edited the final version.

**Data accessibility statement:** Most of the data is publicly available (see methods). The Fish data is available upon request.

**Code accessibility statement:** R functions developed in this paper are available at https://github.com/FloMazel/FD_PD_Max

## Introduction

We are in the midst of a period of heightened biological extinction, with rates several orders of magnitude higher than background rates estimated from the fossil record^1–3^. In addition to having potentially widespread consequences for the functioning of ecosystems and the provisioning of valuable ecosystem services, this situation poses an immense moral challenge^4–8^. Since the extent that resources for conservation actions remain limited, agonizing choices as to which species most warrant attention become necessary^9,10^. To keep humanity’s options open, and our common legacy as rich as possible, it is widely argued that we should seek to maximize the biological diversity of form and function in conservation strategies^6–12^. The biological diversity of form and function can be measured as functional diversity [FD] (see methods). However, in practice, it is challenging to prioritize species on the basis of FD: we have imperfect knowledge about which, and how many traits and functions are important in a given context, how these traits and functions vary among species and across space, and how the importance of traits may change in the future^13^. Many researchers have therefore advocated for a “phylogenetic gambit”; that is, if species traits reflect their shared evolutionary history, then the pattern of that evolutionary history–their phylogeny–should serve as a useful stand-in for unmeasured and unmeasurable traits^9,14,15^. The phylogenetic gambit implies that maximizing phylogenetic diversity (PD), i.e. the breadth of evolutionary history, will ensure that a wide variety of forms and functions are present within a species set^14–17^.

Following this logic, phylogenetic diversity has formed the basis of global conservation schemes, notably the EDGE program^18^ has been used by restoration biologists^19^ and has been widely embraced by researchers across the biodiversity sciences^20–23^. Despite this enthusiasm, the critical question of whether maximizing PD will actually capture more FD than prioritization schemes that ignore phylogeny has, to our knowledge, never been empirically tested^16^. Some studies have discussed^24,25^ and documented the relationship between FD and PD, both at regional^26^ and global scales^20,22^, and many of these studies have shown that maximizing PD does not *maximize* FD. However, such studies do not test the fundamental phylogenetic gambit at the heart of all PD-based conservation strategies: maximizing PD captures *more FD than randomly choosing species*. No one would dispute that the best way to maximize FD is to prioritize FD, but phylogenetic diversity has emerged as prioritization tool because we rarely have sufficient trait data to calculate FD. Here we test whether PD-based conservation passes a much less stringent, but ultimately more fundamental, test: is conserving on the basis of PD better than conserving at random? Worryingly, a recent theoretical study has indeed demonstrated that PD could be a poor surrogate for FD and, in some scenarios, prioritizing species on the basis of PD could actually lead to capture *less* FD than if species were simply selected at random^16^.

This points to the need for empirical tests of whether —within a given species pool— sets of species selected to maximize PD actually contain more FD than sets of species selected without regard to evolutionary relatedness. We clarify what our goals are in testing the utility of PD to capture FD. First, we take as given that maximizing PD is not the overarching goal *per se* of PD-maximization schemes, but rather that a PD maximization strategy is valued for its ability to capture more FD compared to a strategy that ignores phylogeny. Second, it is important to note that we are selecting *species* sets to maximize PD or FD within a region. While this is a simplification, as conservation actions often aim to select sets of *areas* (e.g. in reserve design), the only global phylogenetically-informed conservation initiative is species-centered^18^ (EDGE). Critically, the question we raise has been shown to be distinct from asking whether traits have phylogenetic signal (whether closely related species tend to share similar sets of traits), since PD can be a poor surrogate for FD *even if* traits exhibit phylogenetic signal^16^.

We evaluate the PD^~^FD relationship for different species pools (taxonomic families and geographical assemblages, *i.e*., sets of species co-occurring at a given scale) using a large global dataset including trait, phylogenetic, and geographic range data for 4,616 species of mammals, 9,993 species of birds, and 1,536 species of tropical fish. Specifically, we measure FD as functional richness (see methods) and compute, for any given species pool, an estimate of surrogacy^27,28^ (S_PD_FD_, Figure 1). S_PD_FD_ represents the amount of FD sampled by the set of species chosen to maximize PD, relative to the FD sampled by optimal set of species selected to maximize FD directly, with both components controlled for the expected FD from a random species set of the same size. S_PD_FD_ will be positive if the averaged PD-maximized set contains more FD than the averaged random set, and negative if not. S_PD_FD_ will equal 100% if the PD-maximization strategy is optimal *(i.e*. to maximize FD). We integrate S_PD_FD_ for each species pool across all deciles of species richness but because they are many sets of species that can maximize PD or than can be chosen at random, we computed S_PD_FD_ based on the averaged FD over 1000 PD-maximized sets and 1000 random sets ^16^.

**Figure 1.**
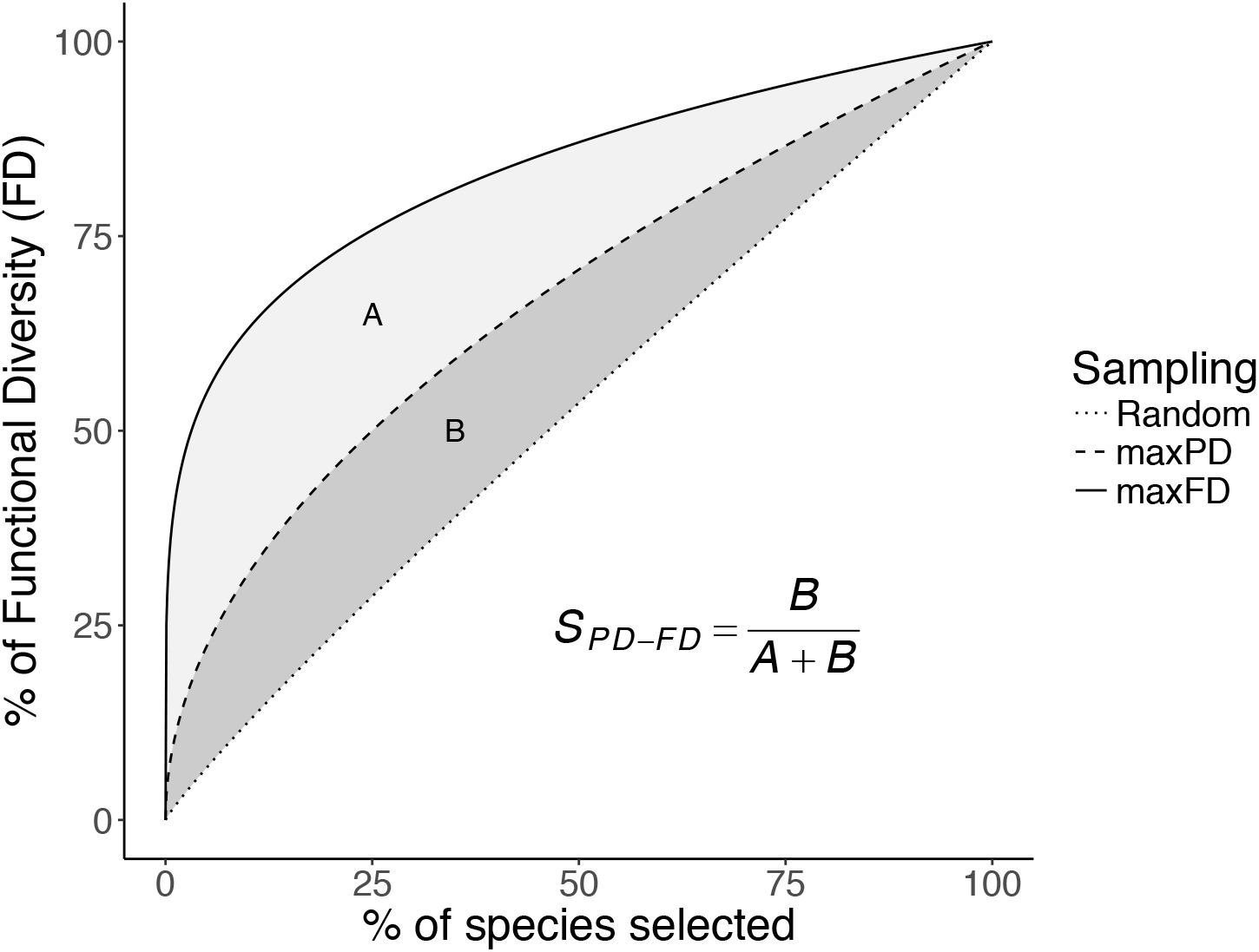
A conceptual approach for evaluating whether PD is a good surrogate for FD. To evaluate if PD is a good surrogate of FD, we measure to what extent a species prioritization strategy that maximize PD captures FD relative to an optimal and a random strategy. To do so, we compare FD accumulation curves (i.e. FD computed for increasing proportion of the species pool considered) across these three different sampling strategies: the random sampling (i.e. rarefaction curve, averaged over 1000 sets), the maxPD (surrogacy, averaged over 1000 sets) sampling (i.e. the sets that maximize PD) and the maxFD (optimal) sampling (i.e. sets that maximize FD, see legends). Then, we measure the surrogacy of PD for FD (SD_PD-FD_) as the area between the random and the maxPD curve (‘A’, see legend) divided by the area between the random and the maxFD curve (‘A+B’, see legend). If SD_PD-FD_ is positive, PD is a good surrogate for FD (the maximum value being 1 where PD is an optimal surrogate) while when SD_PD-FD_ is negative preserving species based on PD is worse than preserving them at random.

## Results

We find that selecting the most phylogenetically diverse sets of species within a given taxonomic family or within a given geographical location (large grid-cells across the globe) captures, on average, 18% more FD than sets of randomly chosen species (i.e. S_PD_FD_ = 18%, SD +/− 6.5% across pools, see Figure 1 and S1-2). Although the surrogacy is generally positive, there was substantial variation across species pools. For example, the surrogacy of PD varies widely from a minimum of −85% to a maximum of 92%, meaning that selecting the most phylogenetically diverse sets of taxa can capture either 85% less (or 92% more) FD than sets of randomly chosen taxa (Fig. 2–3 and Fig. S3-4). However, in 88% of the species pools, choosing sets of species according to PD captured more FD than would be expected at random (*i.e*., surrogacy values > 0 in 88% of the cases, see Fig. 2–3). This suggests that, on average, maximizing PD is a sound strategy to capture FD.

**Figure 2.**
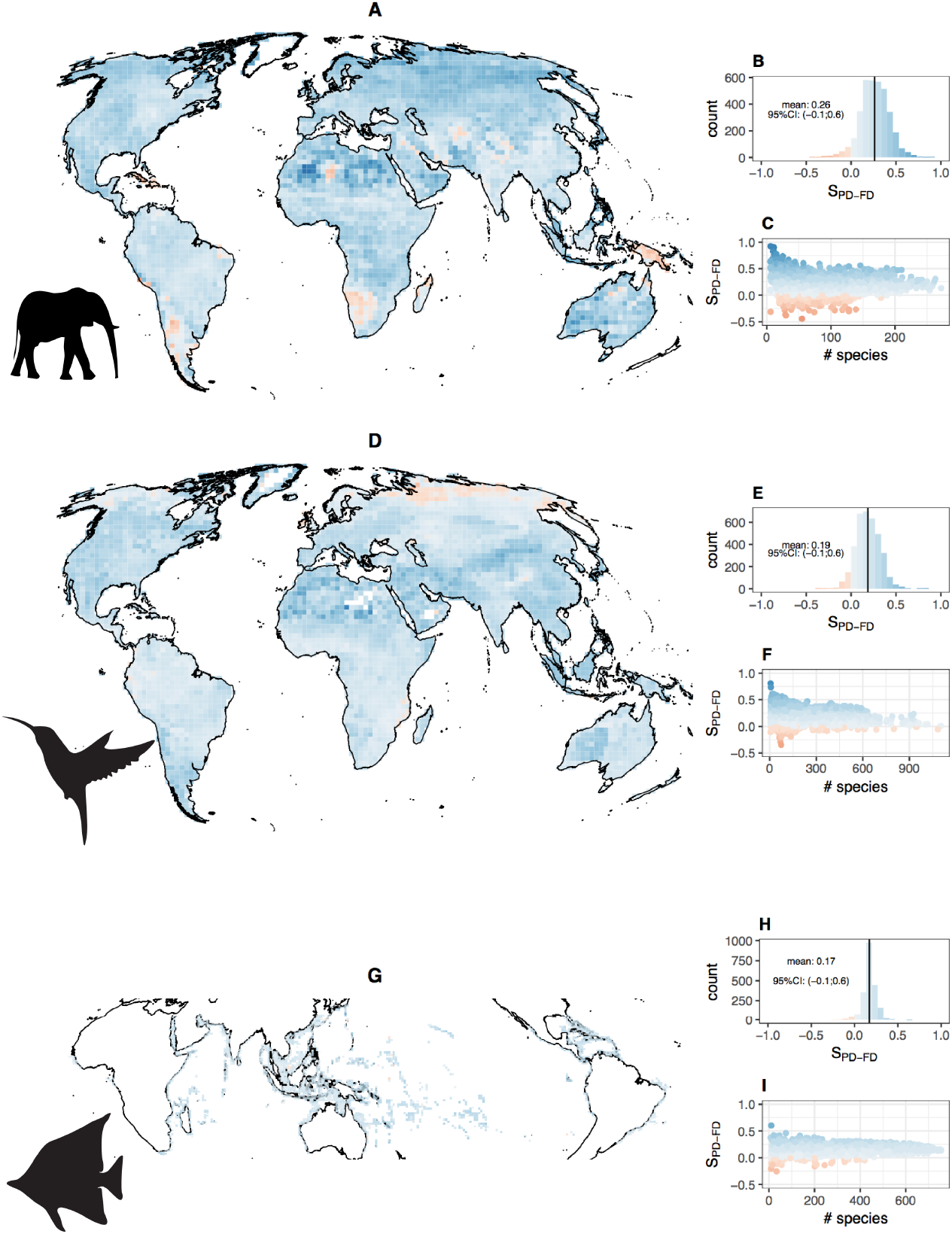
PD is a good surrogate for FD across space. The figure presents the distribution and correlates of S_PD-FD_ for mammals (panels A-C), birds (panels D-F) and tropical fishes (G-I) separately across space. For each of the three groups, the SDpd-fd frequency distribution is presented in top panels (B, E and H) along with its mean (vertical line) and the color code that is common to all panels, with blue indicating positive S_PD-FD_ (maximizing PD captures more FD than random). SDpd-fd geographical distribution is presented in middle panels (A, D, G). Relationships between SDpd-fd and species pool richness are presented in panels C, F and I. In each grid cell, SDpd-fd values are based on the mean over 1000 repetitions of random and PDmax set draw (there is only one maxFD set).

**Figure 3.**
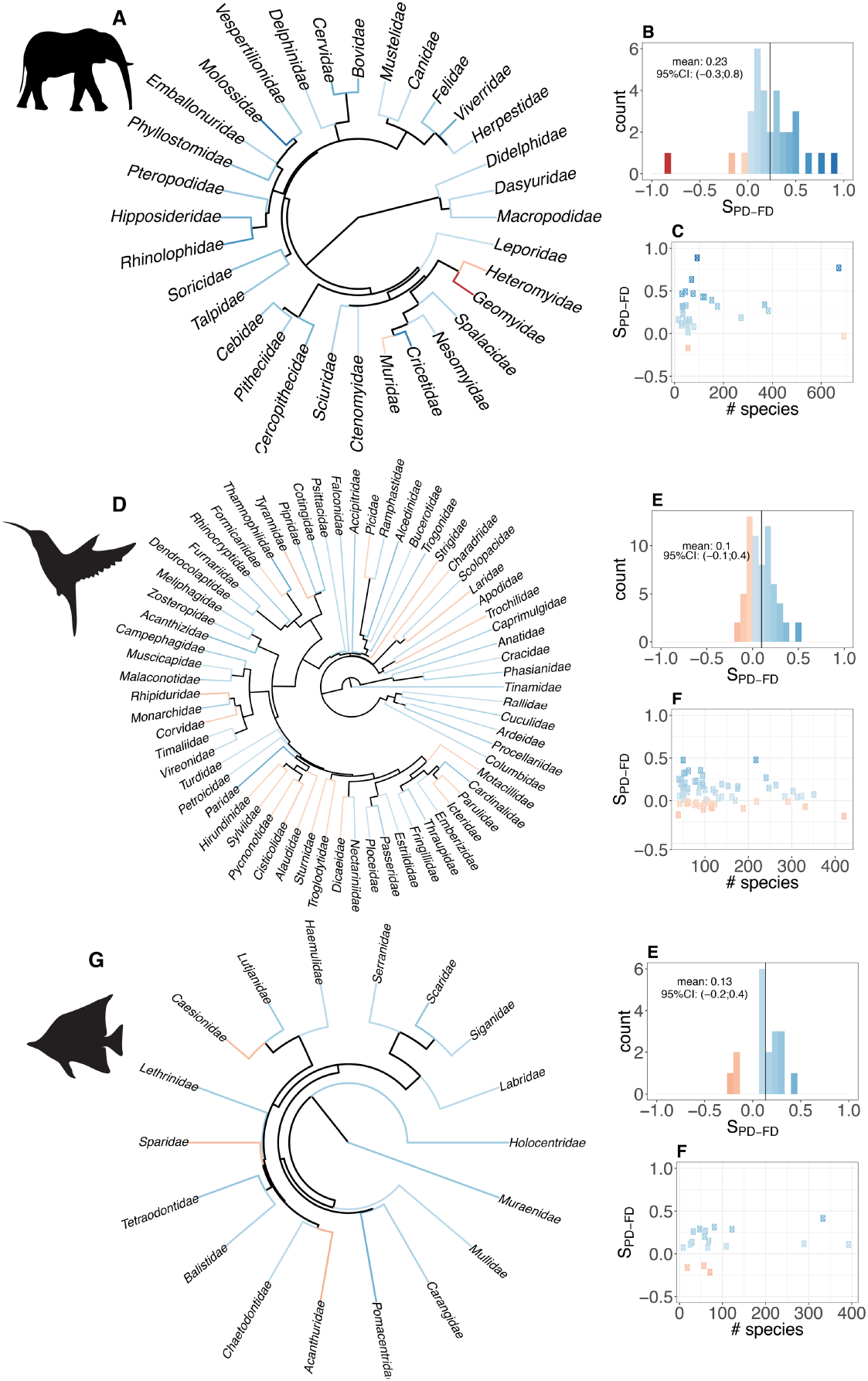
PD is a good surrogate for FD across clades. The figure presents the distribution and correlates of SDpd-fd for mammals (panels A-C) and birds (panels D-F) across families. For each of the two groups, the SDpd-fd frequency distribution is presented (B and E) along with its mean (vertical line). The colour code that is common to all panels. SDpd-fd phylogenetic distribution is presented in panels A and D. Relationships between SDpd-fd and species pool richness are presented in panels C, F and I. For each taxonomic family, SDpd-fd values are based on the mean over 1000 repetitions of random and maxPD set draw (there is only one maxFD set).

However, even if in the majority cases maximizing PD does, on average, better than an averaged random selection, this does not capture the *reliability* of its performance. The PD-maximization and the random selection strategies exhibit variation: simply by chance, random selection of species can capture very high (or, conversely, very low) FD, and the same may be true (to a previously unstudied degree) for PD. The extent of this variation is important: if it is less than the average difference, PD-maximization is a reliable strategy as it will always yield more FD, but if it does not, then PD-maximization could be unreliable for individual conservation interventions. To contrast these two situations, we measured the fraction of times that, within each species pool, the PD-maximization strategy yielded more FD than random selection (see methods). PD-based selection was the best choice in 64% of cases (SD across species pool=9%, see Supplementary Table 1 and Fig. S5), making it the better strategy but not a perfectly reliable one. Thus, while the PD-maximization strategy has a consistent positive effect (i.e. the average PD-maximization strategy yields more FD than the average random strategy), its effect is weak (i.e. the PD-maximization strategy still yields less FD than the random strategy in 36% of the trials within a species pool).

We next explored the drivers of surrogacies values across species pools. Surrogacy of PD appears to weaken as the species pool richness increases (on average, Spearman Rho between absolute surrogacies and species richness = -.15), most clearly seen in the tropics and in species-rich families such as the Muridae (rats, mice and allies) and Columbidae (pigeons and allies) (Fig. 2–3). This is likely because our measure of FD (see Methods) rapidly saturates as the number of selected species increases and species from these large pools harbor high functional redundancy, such that a random prioritization scheme performs relatively well, or at least no worse than other strategies (Fig. S6). In contrast, FD can be greatly increased by prioritization of species using PD from species poor assemblages or clades. This is particularly the case in spatial assemblages containing multiple taxonomic orders, which are both phylogenetically and ecologically divergent from one another. Interestingly, the PD-FD relationship was not consistent across *taxonomic* scale: we found that, in contrast to patterns at the family level, for certain mammalian and avian orders (which are older than the families described above), using PD to select species is much worse for capturing FD than choosing species at random (see, for example, the Afrosoricidae, Chiroptera, and Charadriiformes in Fig. S7).

We then explored whether we can explain this variability within- and between-datasets, and in particular, why for some assemblages/clades, a PD-prioritization strategy fails to capture more FD than random choice. It is often implicitly assumed that phylogenetic signal (i.e. the degree to which closely related species tend to harbor similar sets of traits) can be used to evaluate the effectiveness of PD as a surrogate for FD ^5,15–17^ Surprisingly perhaps, the value of PD as a surrogate for FD was only weakly correlated with the phylogenetic signal of the underlying traits (Fig. S8-9, on average Spearman Rho = 0.17). Similarly, tree imbalance, which is known to affect surrogacy in simulations ^16^, did not explain surrogacy in these empirical data (Fig. S8-9).

For mammals, regions where PD did worse than random were located in the Sahara, south western Patagonia, southern Africa including parts of Madagascar, and New Guinea (Figure 2). These latter two in particular are of concern since they are global conservation priorities on the basis of species endemism and habitat loss. We suggest two historical reasons for such idiosyncratic poor performance of PD. First, there is a tendency for a large carnivore species, either a top predator *(e.g.*, cheetahs in the Sahara or foxes in Patagonia) or a large scavenger (e.g., the hyena in South Africa) to co-occur with a close relative with distinct traits in these areas (e.g., a desert cat with the cheetah or the aardwolf with the hyena, see Fig. S10). Only one of these closely-related species will tend to be selected under prioritization schemes that maximize PD, thus reducing the volume of the convex hull on average when the functionally distinct one is not selected (the large predator or scavenger). This seems also to drive the low surrogacy of PD in Charadriiformes (especially *Larus* and *Sterna;* see Figure S10). Second, lineages in which traits evolve very slowly will contribute little to FD, even over long periods of time (branch lengths) that contribute greatly to PD. For example, in New Guinea many co-occurring bats with similar traits diverged long ago, such that they are always selected in the PD maximizing set, but do not add much to the convex hull, resulting in a poor surrogacy of PD for FD. Such strong ecological niche conservatism is common in mammals ^29^, e.g. in the Geomyidae: two basal branches of the Geomyidae tree harbor very similar traits (species descending from these branches are actually grouped in the same genus *Thomomys)* while being distantly related in the phylogenies we used (Fig. S10). As such, they will be selected in all PD maximizing sets, but will not contribute greatly to FD.

## Discussion

Maximizing PD in conservation decisions is now commonplace in the academic world^20–22,30–33^ and is also starting to be used in real-world conservation prioritizations, for example with the EDGE program^18^. To the best of our knowledge, there are no clear direct ecosystem function or health benefits that phylogenetic branch lengths provide. Rather, high PD is perceived as valuable because it is assumed to be a good proxy for high diversity of traits or “features”^14^ (referred as to high functional diversity in this paper, FD), a hypothesis that we name the “phylogenetic gambit”. High FD might be valuable for a number of reasons, for example ecosystem functioning, ecosystem services, future “options values”^14,15^ or “evolutionary potential”^15,34^. The utility of PD for conservation stems from the fact that calculating PD is relatively fast and cheap, often making it an easier way to prioritize species or areas than FD. Indeed, we have imperfect knowledge about which, and how many, traits and functions are important in a given context, how these traits and functions vary among species and across space, and how the importance of traits may change in the future^13^. Yet, even if convenient, maximizing PD can only be an effective and realistic conservation strategy to conserve FD if the phylogenetic gambit holds and maximizing PD yields more FD than a strategy that ignores phylogeny. If maximizing PD yields *less* FD than a random strategy (i.e., the gambit fails), then researchers and conservationists should reconsider whether maximizing PD as a useful conservation strategy. A large body of literature has shown that maximizing PD does not maximize FD empirically^20–22,30^ or even in simple theoretical cases^16^, but such work does not test the phylogenetic gambit of whether PD prioritization captures more FD than *random* selection (which has not, to our knowledge, been tested)^16^. Here we have shown that the phylogenetic gambit holds: that PD *is* an effective conservation metric to capture FD. Yet we also show that it remains something of a gambit: PD is good ‘on average’, but there is still some risk associated with taking it.

We found that prioritizing the most phylogenetically diverse set of taxa in a region or clade will result in an average gain of 18% functional diversity relative to applying the same conservation effort without considering phylogeny, but this gain will decrease as species richness increases. In opposition to what has previously been implicitly assumed^15,16^, we find weak empirical evidence that the presence of phylogenetic signal in traits predicts whether PD-based conservation will prioritize FD. Our result suggests that PD is a reasonable conservation prioritization strategy, especially in species-poor clades or regions, or in the absence of meaningful data on functional traits. However, we note three important caveats to the use of this strategy. First, 18% extra FD may not always be a useful conservation target. It is currently unknown whether this added 18% of FD can actually be of enough conservation value. Second, in cases of either recent trait divergence or, alternatively, very strong trait conservatism, a PD prioritization scheme can capture less FD than a random scheme. Evolutionary biologists commonly focus on ‘unusual’ clades with rapid divergences (e.g., cichlids); we show here that divergence does not have to be that spectacular (e.g., African carnivores) to alter the PD-FD relationship. Third, we found that while this strategy, on average, captures FD well, it is also somewhat unreliable, and 36% of the time will not capture more FD than random choice. This means that while the PD gambit can be a bet worth taking, it is still a bet with associated risk, not a sure thing.

Our objective in this paper is to test the phylogenetic gambit using empirical datasets. This means that we do not aim to provide a coherent prioritization strategy^35^, or ready-to use conservation guidelines. Indeed, we simplistically and implicitly assume that chosen species will either be saved or go extinct, and we have not linked our various scenarios to any particular policy position or conservation objective other than maximizing FD within a phylogenetic clade or region^28,31^. In reality, conservation decisions reflect the interplay of social, economic, political, and scientific priorities, and do not necessarily result in the saving of target species (and therefore of their associated FD or PD). While our study is thus not directly applicable, the test we are conducting is actually critical to validate (or invalidate) the use of PD in conservation as a whole. While it is not clear whether our results would generalize to other taxa (although we hope that others will extend our work and test the phylogenetic gambit in other systems), we do feel it is important to consider the uncertainty that has been introduced into our analysis as a result of uncertainty associated with the spatial scale of our analysis, our phylogenetic data, and our choice of trait and measurement of FD.

The scale of conservation activities can vary, from the global scale of the hotspots approach to local protected areas within a single country, but, unfortunately, the connection between these scales remains unclear. For example, if the motivation for protecting FD is to maintain community-driven ecosystem functions and services ^6,36,37^, the value of a regional or global focus may be questionable^38^, and studies are increasingly focusing on local scales^6^. Ecologists are refining and improving our understanding of how local assemblages assemble within a regional context^39^, and while the concept of the ‘regional pool’ of species is increasingly being viewed as a simplification, it is unlikely that regional- and local-scale patterns are totally disconnected. We emphasize that our results are relatively robust to variation in spatial scale (see Fig. S3), but we acknowledge that future studies should test the phylogenetic gambit at more local scale as well.

The set of species that maximize PD obviously rely on the phylogenetic hypothesis used. No hypothesis is perfect or without uncertainty, and these phylogenetic uncertainties could in turn impact the composition of the set of species that maximize PD and hence the surrogacy values we compute. In this study, we explicitly took into account these uncertainties by using 100 different trees^40,41^. The explicit propagation of this phylogenetic uncertainty through into our results may underlie some of the uncertainty (‘risk’) of our result, and we suggest it is important that future studies explicitly take into account phylogenetic uncertainty when testing the phylogenetic gambit.

The motivator of our test of the surrogacy value of PD for FD is the fact that ecologically-relevant trait data is in short supply, especially for rare and data-deficient species. Indeed, if it were not for this relative paucity of data, we could simply prioritize species based on their unique contribution to FD directly. Although there have been massive and well-funded efforts to collect and curate trait data from across the Tree of Life^42–44^, we are still far from having comprehensive coverage. Furthermore, despite recent progress^45^, it is still not fully understood which traits are most relevant for responses to environmental change, or that contribute most to certain ecosystem functions and services, and how these vary among systems. Our analysis suffers from a similar data limitation. We chose these traits because they are frequently collected in ecological studies, not because we know they are ecologically important. Our assumption is that their phylogenetic distribution is typical of those traits that are most desirable for the purpose of conservation and that our primary results are therefore widely applicable. While we did test the robustness of our results to the variation of trait information retained to compute FD (Figure S1), it is true that, overall, we used a rather limited set of traits. We acknowledge that it is possible that many other potential valuable traits are not captured by our measure of FD. One of the ideas behind the use of PD is that phylogeny might account for these for unmeasured and unmeasurable traits^9,14,15^, however, as this hypothesis is not testable (we do not have these traits), it seems risky to assume it is true. Our objective here is to test the phylogenetic gambit given the limited set of traits that we have: we consider that carrying out our imperfect test is more informative than not carrying any test at all.

In conclusion, we found that maximizing PD results in an average gain of 18% of FD relative to random choice. However, this average gain hides the fact that in over 1/3 of the comparisons, maximum PD sets contain less FD than randomly chosen sets of species. These results suggest that, while maximizing PD can help capture FD, it represents a risky strategy. If maximizing PD is a risky strategy, then, should we abandon the use of PD in conservation? We believe that before such dramatic decision, our test should be repeated across space, traits and taxa, in order to narrow the uncertainties of our results. This is why we now urge others to expand our simple phylogenetic gambit test to other clades and other traits in order to test the generality of our findings. We hope that our study will stimulate the production of numerous tests to finally rigorously assess the usefulness of PD in conservation.

## Acknowledgements

This paper is a joint effort of the sCAP working group funded by the Synthesis Centre of the German Centre for Integrative Biodiversity Research Halle-Jena-Leipzig and the Canadian Institute for Ecology and Evolution. FM was supported by an NSERC Accelerator Grant to AOM and a Banting Post-doctoral fellowship. GDR thanks Mike Steel. We thank Risa Sergeant at the University of Ottawa for hospitality, the Crawford Lab for Evolutionary Studies at SFU for discussion.

## Material and Methods

We use two classes of data to address the question of whether choosing sets of species according to PD captures the underlying trait diversity (as measured with FD) well. First, we used taxonomic groups (clades) of species as our unit of analysis (‘species pool’ hereafter) and, second, we investigated broad assemblages found across the globe. The former is more explicitly evolutionary, ensuring that our results are not driven by well-established relationships across large taxonomic groups (e.g., monotremes are distinct from placental mammals) and the latter is likely more relevant to actual conservation practice.

### 1. Data

We use distribution data to delineate geographical assemblage species pool and taxonomy to delineate clade-based species pools (namely families and orders).

#### Distribution data

For mammals, we used the distribution maps provided by the Mammal Red List Assessment (http://www.iucnredlist.org/) for 4,616 species. For birds, full (breeding and wintering) and breeding ranges distribution maps were extracted from BirdLife (http://www.birdlife.org/) for 9,993 species. The best resolution at which these maps should be used is still under discussion in the literature, so we decided to use the 40 000km^2^ resolution (200×200km gird cell at the equator) that is commonly used at global scale ^46,47^. The total number of grid cells was 3,646. Domestic and aquatic mammals were excluded from the analysis. In order to make sure our results were not driven by the important trait difference between volant and non volant mammals, we repeated our results excluding bats. For birds, we repeated our analysis using the full ranges. Finally, we evaluated the robustness of our result to the spatial resolution considered by repeating our analysis at a resolution of 100×100km (number of cells was 13,330) for birds and mammals; we present these results in the supplementary materials, as they are qualitatively identical to those conducted at 200×200km (Fig. S1). For fishes, we used a database of 1536 species, for which we had distribution data, phylogenetic and functional data. Distribution data were extracted from a global-scale distribution database ^48^. Species composition was then extracted from grid cells of 5°x5°, corresponding to approximately 555×555 km at the equator ^49^. This grain size of the grid was chosen because it represents a good compromise between the desired resolution and the geographical density of information.

#### Phylogenies

In order to prioritize species to maximize PD, phylogenies of each species pool are needed. We used the first 100 published calibrated ultrametric trees of Jetz et al. (2012) for birds and Faurby and Svenning (2015) for mammals. By repeating our analyses across a posterior distribution of phylogenetic hypotheses, we control and account for phylogenetic uncertainty. For tropical reef fishes, we built a phylogeny for 18 families (i.e. Labridae, Scaridae, Pomacentridae, Chaetodontidae, Acanthuridae, Haemulidae, Balistidae, Carangidae, Serranidae, Lutjanidae, Sparidae, Caesionidae, Holocentridae, Mullidae, Muraenidae, Tetraodontidae, Lethrinidae and Siganidae) by pruning a dated molecular phylogenetic tree for 7,822 extant fish species ^49^. These families were selected as the most representative tropical reef fish families, that is, they are abundant and speciose on tropical reefs. We grafted missing species on the pruned phylogenetic tree (circa 50% among the 1536 studied species) based on published phylogenies for these families, supplemented by taxonomic information from fish identification guides and FishBase ^49,50^. We recorded, for each of these trees, a measure of imbalance (as measured by **β** ^51^) and ‘tipiness’ (as measured by Gamma ^52^). For both mammals and birds, we chose to group species in families and orders. We used these groupings when calculating the purely phylogenetic, clade-based analyses (to address question 1), but not within the spatial, assemblage-based analyses (question 2). For the taxonomic analysis of mammal families, we removed two families (Dipodidae and Echimyidae) because of their very poor phylogenetic resolution (i.e. polytomies for an important number of species).

#### Traits

For birds and mammals, four traits (diet, (log transformed) body mass, activity cycle, and foraging height) were extracted from Elton Traits1.0 ^44^. These traits are generally assumed to appropriately represent Eltonian niche dimensions within an assemblage or clade of mammals or birds ^53,54^. For fishes, we used a previously published database ^12^. We used 6 categorical traits: size, mobility, period of activity, schooling, vertical position in the water column, and diet (for a full description of the dataset, see Mouillot *et* al. 2014). These traits have already been used to investigate community assembly rules ^55^ and to seek vulnerable fish functions ^11^. For each clade and assemblage, we used the raw trait (only body mass was log-transformed and rescaled by the clade/assemblage range of body masses) values to compute distance between species using Gower distance [19] and use PCoA to summarize the trait space in few dimensions. We retained the numbers of PCoA axes necessary to represent 70% of the total initial variability (using a 80% threshold did not quantitatively change our conclusions, see Fig. S1). We also recorded phylogenetic signal for each PCoA axis using Blomberg’s K ^56^.

### 2. Approach

Our aim was to evaluate, across a wide range of clades and regions, the ability of PD-informed prioritization scheme to capture FD in comparison with two other prioritization schemes: selecting species to directly maximize FD (‘maxFD’ hereafter) and selecting species randomly (Figure 1). Our premise was that we often do not know or have not measured the traits that are most relevant for ecosystem function and services such that maximizing FD is not generally feasible. By focusing on a subset of traits and assuming that they are representative of ecologically relevant traits, we were able to get an estimate of how well PD does compared to the best we could possibly do. We used performance relative to choosing on the basis of FD as an upper-limit to the performance of PD as a surrogate for FD, and used random species selection as a lower benchmark.

#### Random prioritization scheme

For each pool (i.e. each clade and each geographical assemblage) and each number of selected species (10, 20, 30, 40, 50, 60, 70, 80, 90, and 100% of the total pool), 1000 random sets of species were produced, from which the average FD was recorded.

#### Prioritization scheme maximizing PD (maxPD)

While there are many, overlapping metrics for measuring the evolutionary history encompassed by a set of species ^15,57^, the most common is the sum of all branch lengths (often in units of time) connecting a set of species to a common root ^14^, called Phylogenetic Diversity (PD). This is the metric whose maximization has most commonly been proposed as a conservation prioritization metric ^14,34,58^, and as a measure of phylogenetic ‘richness’ it most naturally maps onto our chosen FD metric ^57^. We used the greedy algorithm proposed by Bordewich *et* al. (2008) to find our maxPD set of species S. For a given tree there are likely multiple, and possibly very many, sets of species with the same PD as S. As a consequence, we produced, for each pool, each number of selected species, and each alternative phylogenetic trees, 10 maxPD sets of species. We then averaged the FD of these sets across our 100 phylogenetic trees, so that each value is an average of 1000 sets (10 sets for each of the 100 trees).

#### Prioritization scheme maximizing FD (maxFD)

Functional diversity was estimated using a functional richness index (FRic; Cornwell *et* al. 2006; Villéger *et* al. 2008; Pavoine & Bonsall 2011). The FRic index relies on a multidimensional Euclidean space, where the axes are traits (or factorial axes from a principal coordinates analysis (PCoA) computed using these traits) along which species are placed according to their trait values. This index measures the volume of trait space occupied by a given species assemblage by calculating the convex hull volume ^62^, defined by the species at the vertices of the functional space, that encompasses the entire trait space filled by all species in this assemblage. In a single dimension, this simply equals the range of values ^62^. This broadly used metric in ecology is set monotonic with species richness, a property generally assumed desirable in conservation whereby the addition of a new species can never decrease the metric’s value ^63^. FD measures the total amount of variation in trait values, making it conceptually comparable to PD ^57^. We used the FRic index instead of the FD index based on a functional dendrogram (Petchey & Gaston, 2006) since recent studies showed that the FD index may lead to biased assessments of functional diversity and inaccurate ecological conclusions ^64^. The most straightforward way to obtain the maximal FD for *n* species is to compute FD for all possible combinations of *n* species and simply record the greatest value (the brute force approach). However, this is not feasible in practice as the numbers of combinations of selected species was too high (e.g., 10^71^ possible sets for all mammal assemblages). To rapidly and efficiently find the set of species that aim to maximize FD, we developed a novel (at least in ecology) greedy algorithm. In brief, our approach iteratively (starting with two species) select the species that is the furthest from the centroid of the already selected set. To avoid selecting two species that are far from the centroid but close to each other, we penalized the distance to the centroid by the distance to the closest neighbour in the already selected set. Here we present in details the greedy algorithm we used to find the set of species that maximize FD:

Step 1. Select the two species with the highest trait distance
Step 2. Compute the centroid of these two selected species
Step 3. Compute distances between species not in the set and this ‘set centroid’.
Step 4. Penalize these distances by adding the following factor f (Eq. 1)

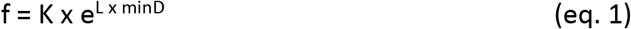

with K and L being penalizing factors and minD the distance between a given candidate species and the nearest species already in the selected set.
Step 5. Select the species that maximized the penalized distance
Step 6. Go back to step one with this new set of species until the desired number of species is reached.

To avoid arbitrarily setting the penalizing parameters, we tested 1000 pairs of parameters drawn from a truncated normal distribution (mean=1, sd=.5) and retained the parameter pairs that yielded the maximal FD.

In tests of subsets of the data for which finding the true maxFD was feasible, we found our approach to adequately approximate the true maxFD and to produce a very good approximation of the true degree of PD’s surrogacy for FD (Fig. S2).

#### Measuring performance and surrogacy of prioritization schemes

We use a common approach^27,28^ to quantify the extent to which a given surrogate (here, the maxPD choice) reaches a certain objective (here, maximize FD). Species from a given pool (i.e., for each dataset (clade and assemblages) independently,) were prioritized and selected according to (1) the objective, i.e. maximize FD, producing the ‘optimal curve’ (maxFD curve in Figure 1), (2) the surrogate i.e. maximize PD, producing the ‘surrogate curve’ (maxPD curve in Figure 1) and (3) at random (random curve in Figure 1), i.e. producing the ‘random curve’ (Figure 1). To compute a ‘surrogacy’ estimate of PD (S_PD-FD_), we compare the position of the surrogate curve (1) to the random curve (2) relative to the optimal curve (2) (Figure 1 and Eq. 2) across the deciles of species richness of the pool (given as an interval 0-1):

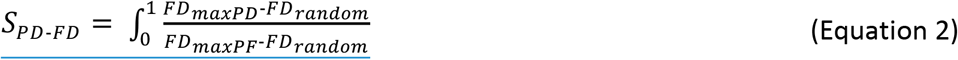

This surrogacy metric is at 100% when the surrogate perfectly meets the objective (i.e., the maxFD and maxPD curves are identical and the max PD set is the maxFD set), 0% when the surrogate is not better than randomly chosen sets of species (i.e., the random and maxPD curves are identical) and is negative if the surrogate choice is worse than random (i.e., the maxPD curve is below the random curve). Correlates of S_PD-FD_ were evaluated using Spearman correlations.

Apart from focusing on average tendencies, we quantified the variability of the FD yielded by the PD—maximized selection strategy and the random selection strategy within each species pools. To do so, we compute, for each species pool and for each % of selected species independently, the number of cases where FDrandom>FDmaxPD across the 1000 random *1000 maxPD sets combinations (i.e. 10^6^ comparisons). We then averaged theses number across % of selected species and report statistics across datasets (Supp. Table 1).

